# Dynamics of Changes in Cadmium Levels in Environmental Objects and its Impact on the Bio-Elemental Composition of Living Organisms

**DOI:** 10.1101/2023.01.22.525090

**Authors:** Larysa Nechytailo, Svitlana Danyliv, Liliia Kuras, Svitlana Shkurashivska, Aleksandra Buchko

## Abstract

As a result of intensive anthropogenic impact in the biosphere there is a rapid process of accumulation of heavy metal salts. They have led to the aggravation of problems associated with the pollution of ecosystems and basic food products of plant and animal origin. Environmental pollution by these compounds is caused by their persistence in environmental objects, migration ability, accumulation by plants. This contributes to their accumulation in the human environment. A number of studies have shown that heavy metals have mutagenic, toxic effects and affect the intensity of biochemical processes. Therefore, the presence of heavy metals in the environment is extremely undesirable. Moreover, the ecological state of the environment is directly related to changes in the human internal environment. Deficiency or excess of certain bioelements in soils and drinking water or non-compliance with its stable chemical composition causes the development of dysmicroelementosis. The ecological situation of the Carpathian region is closely related to the state of soils and water resources. In this regard, it is advisable to study and control the level of cadmium compounds in the environment of the region. The study of the effect of cadmium intoxication on the macro- and microelement composition of the brain and myocardium of experimental animals is also worthwhile.

**Materials and methods:** Soils and drinking water of the plain, foothill and mountainous zones of the region, as well as organs and tissues of experimental animals served as the object of research. Cadmium levels in drinking water and myocardial tissues and brain of experimental animals have been measured by atomic absorption spectroscopy.

**Results and discussion:** The study of soils in the Prykarpattia region has revealed an increase in the toxic element cadmium. Its content is 1.1-1.5 times higher than background levels. The analysis of drinking water allowed to establish that a significant number of people living in the plain and foothill zone of the region consume water with a high content of cadmium. The main stages of cadmium intake and accumulation in plants have been analyzed.

Significant disorders in the body of experimental animals under conditions of excessive intake of cadmium compounds have been revealed. It was accompanied by the accumulation of cadmium in the myocardium and brain, on the background of redistribution of vital macronutrients calcium and magnesium along with micronutrients copper and zinc. Thus, excessive intake of cadmium salts causes the development of dysmicroelementosis, which is accompanied by a violation of the homeostasis of a living organism. It is suggested to conduct continuous monitoring of the level of toxicants in the ecosystem as an integral component of environmental monitoring.

## Introduction

Due to the intensive industrial development, the increase in the number of vehicles, the widespread use of agrochemicals and mineral fertilizers, the biosphere is experiencing a rapid process of accumulation of heavy metal salts. This has led to the aggravation of problems associated with the pollution of ecosystems and basic food products of plant and animal origin.

Heavy metals (HM) are one of the largest toxicants of anthropogenic genesis. Soil occupies a special place in the biosphere among all geophysical environments. It provides biological productivity and is subject to the greatest anthropogenic impact, as it is one of the links in the migration of pollutants [1]. One of the most harmful toxicants is cadmium. The main sources of soil contamination with cadmium are identified as metal processing industry wastes, irrigation of agricultural land with wastewater and application of its sediments to the soil as fertilizers and the use of mineral and organic fertilizers, industrial emissions into the atmosphere from heavy industry, fuel combustion products [2].

Among the measures that should ensure the protection of human health and the environment from anthropogenic metals, an important place is occupied by control over the level of metal content in soil, water, human and animal organisms.

## Literature review

Heavy metals, unlike organic pollutants, do not decompose, but change from one form to another. In particular, they are part of salts, oxides and organometallic compounds, so this is what determines the danger of their release into the environment [3]. Phosphorus and complex fertilizers play a significant role in soil pollution [4, 5]. It is established that with excessive application of superphosphate to the soil, the content of cadmium in vegetable products increases by 4 times compared to the control [6]. With mineral fertilizers, 3-4 g/ha of cadmium is annually applied to the soil, sometimes this value can reach 10 g/ha [7]. Uncontrolled application of fertilizers can lead to soil contamination, from where these toxicants, due to the processes of migration and translocation, enter surface and groundwater, and with them into the organisms of plants and animals. The high level of anthropogenic soil pollution causes the increased content of toxicants in water bodies. Heavy metals coming from the water source can accumulate in plants to levels that cause phytotoxicity [8]. They are resistant to degradation in natural conditions, can accumulate in aquatic flora and fauna [9], enter the food chain and adversely affect the physiological functions of human organs and tissues. The presence of cadmium in drinking water, even in small amounts, requires special attention, as it is characterized by high toxicity and cumulative, carcinogenic and mutagenic properties [10,11]. A feature of the harmful effects of cadmium is its rapid absorption by the body and slow excretion [12]. From literature sources [13, 14] it is known that cadmium accumulates in a number of organs and tissues, mainly in the kidneys, liver, bones, lungs, disrupts metabolic processes and physiological functions, is an antagonist of a number of vital macro- and microelements. The toxic effect of cadmium is to impair oxygen transport to tissues, resulting in inhibition of tissue respiration by binding sulfhydryl groups of enzymes [14, 15]. In this regard, it is advisable to study the brain, as the most sensitive organ to oxygen deficiency, and the myocardium. It is an important organ that stimulates blood transport through the vessels of the body, providing nutrition and respiration of tissues of living organisms. At the same time, it is known that cadmium damage causes the development of hypertension, coronary heart disease, bronchitis, pharyngitis and other respiratory diseases [16,17]. However, the bioelemental composition of the myocardium and brain under conditions of cadmium chloride intake remains poorly studied. It is important for understanding their influence on the course of metabolic processes in living organisms.

**The aim of the research** is to carry out a comparative analysis of the level of cadmium compounds in soils and water bodies of different geographical zones of the Prykarpattia region, as well as to study the effect of cadmium intoxication on the macro- and microelement composition of the brain and myocardium in experimental animals.

## Materials and methods

Soils and drinking water of the plain, foothill and mountain zones of the region, as well as experimental animals were the object of research. The selection of soil and water study sites was carried out taking into account the altitude, seasonal changes, peculiarities of water supply sources and relative distance from the pollution sources. Soil and water samples were taken once in spring and autumn. The methodology of soil and water sampling, their preparation for analysis, transportation and storage was carried out in accordance the State Standardization System [18, 19]. The study of samples for cadmium content was carried out according to DSTU 4770.3:2007 of the State Standardization System. Soil quality. Determination of the content of mobile cadmium compounds in soil in buffered ammonium acetate extract with pH 4.8 by atomic absorption spectroscopy [20]. Cadmium levels in drinking water and myocardial tissues and brain of experimental animals were determined with the help of atomic absorption spectroscopy [21].

Experimental studies were conducted on laboratory animals - white outbred sexually mature male rats weighing 180-220 g. The maintenance of animals, their nutrition and manipulations were carried out in compliance with ethical and legal norms and requirements for scientific and biochemical research: Annex 4 to the “Rules for conducting work using experimental animals”, approved by the order of the Ministry of Health of Ukraine № 755 of 12.08.1997. “On measures to further improve the organization of forms of work using experimental animals” and the provisions of the “General principles of experiments on animals”, adopted by the First National Congress on Bioethics (Kyiv, 2001); according to the Law of Ukraine № 3447-IV “On the protection of animals from cruel treatment” of 21.02.2006; European convention for the protection of vertebrate animals used for experimental and other scientific purpose: Council of Europe 18.03.1986. – Strasbourg [22, 23].

Models of toxic damage to animals were intoxication with cadmium chloride. It was injected intramuscularly at a dose of 1.2 mg/kg (in saline) of animal body weight (1/10 LD_50_) [24] once a day for 10 days. Control animals were simultaneously injected with an appropriate amount of 0.9% sodium chloride solution. Brain and myocardial homogenates were used for the study [25]. The obtained results were subjected to statistical analysis according to the generally accepted method [26] using Student’s t-test (Statistica 8).

## Results

The ecological situation of the Prykarpattia region is closely related to the state of soils and water resources. In this regard, it is advisable to study and control the level of cadmium compounds in the region’s environmental objects.

The study of the content of toxic element cadmium in the soils of the Prykarpattia region allowed to establish that its level in the soils of the plain zone and foothill zone significantly increases in autumn and is respectively: 0.19 mg/kg and 0.10 mg/kg. In the soils of the mountainous zone, the cadmium content was at the level of 0.06-0.16 mg/kg. The tendency to increase the level of this toxicant was observed in autumn and spring (Figure 1).

**Figure 1.**
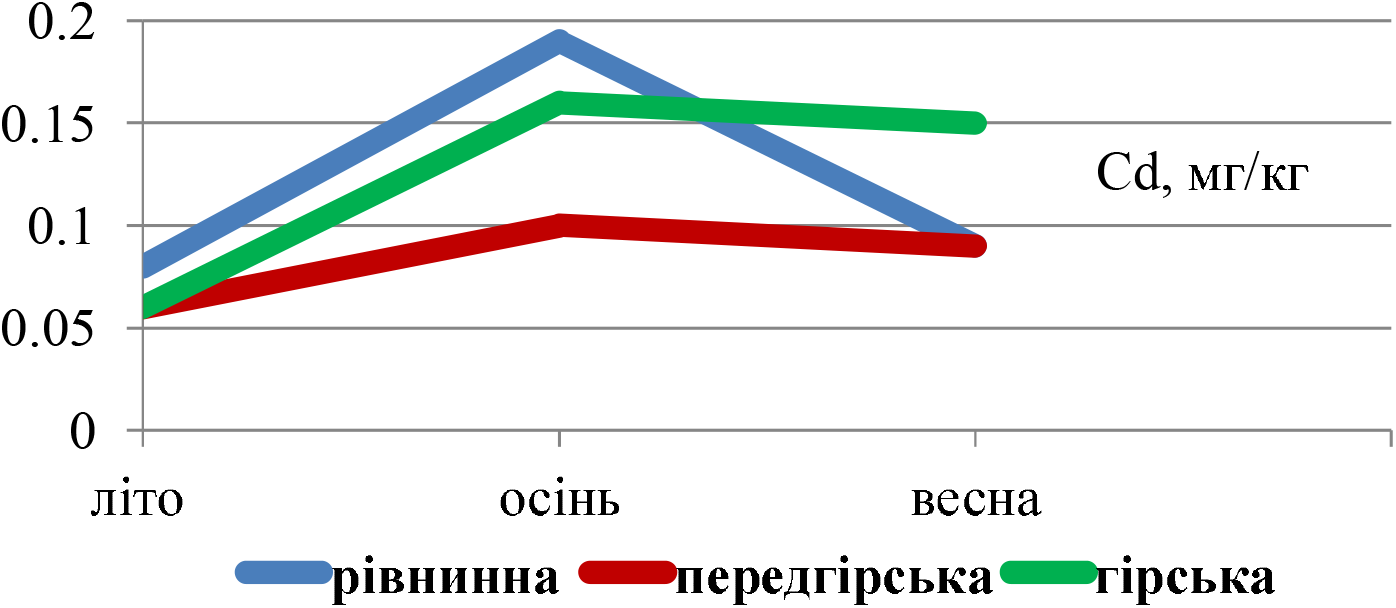
Dynamics of seasonal changes of cadmium level in soils Note: MPC for cadmium - 0.7 mg/kg, background value - 0.136 mg/kg

In order to assess the degree of soil contamination with cadmium, the actual concentrations of the element in the studied samples were compared with the maximum permitted concentration (MPC) and the background indicator. The analysis of the obtained data indicates that the level of cadmium in the soils of the Prykarpattia region did not exceed the MPC. However, attention should be paid to the increase of cadmium above the background level by 1.1-1.5 times in autumn in mountainous and plain zones (Figure 1).

It is known from the literature [27] that due to the migration ability, heavy metals are concentrated in water bodies. Therefore, it is important to study and control the level of cadmium in drinking water in different geographical zones of the Prykarpattia region.

The study of the level of toxic element cadmium in the drinking water of the Prykarpattia region allowed it to establish that its content exceeds the MPC in the areas of the plain zone by 1.6-2 times during all periods of observation. Analysis of the cadmium content in drinking water from the sources of the foothill zone showed a significant increase of this trace element by 1.3-1.9 times above the MPC, in the water of the mountainous zone it did not exceed the maximum permitted values. Thus, in the drinking water of the plain and foothill zones, there is an intensive increase of cadmium in quantities that significantly exceed the maximum permitted level, which poses a real threat to the existence of living organisms.

Among heavy metals, cadmium is extremely toxic to plants and is considered one of the most dangerous elements entering the environment, even at low concentrations [28]. Once in the soil, cadmium is absorbed by the root system of plants, to a lesser extent - through leaves [29], accumulates in them and can enter the body of animals and humans through food chains (Figure 2).

**Figure 2.**
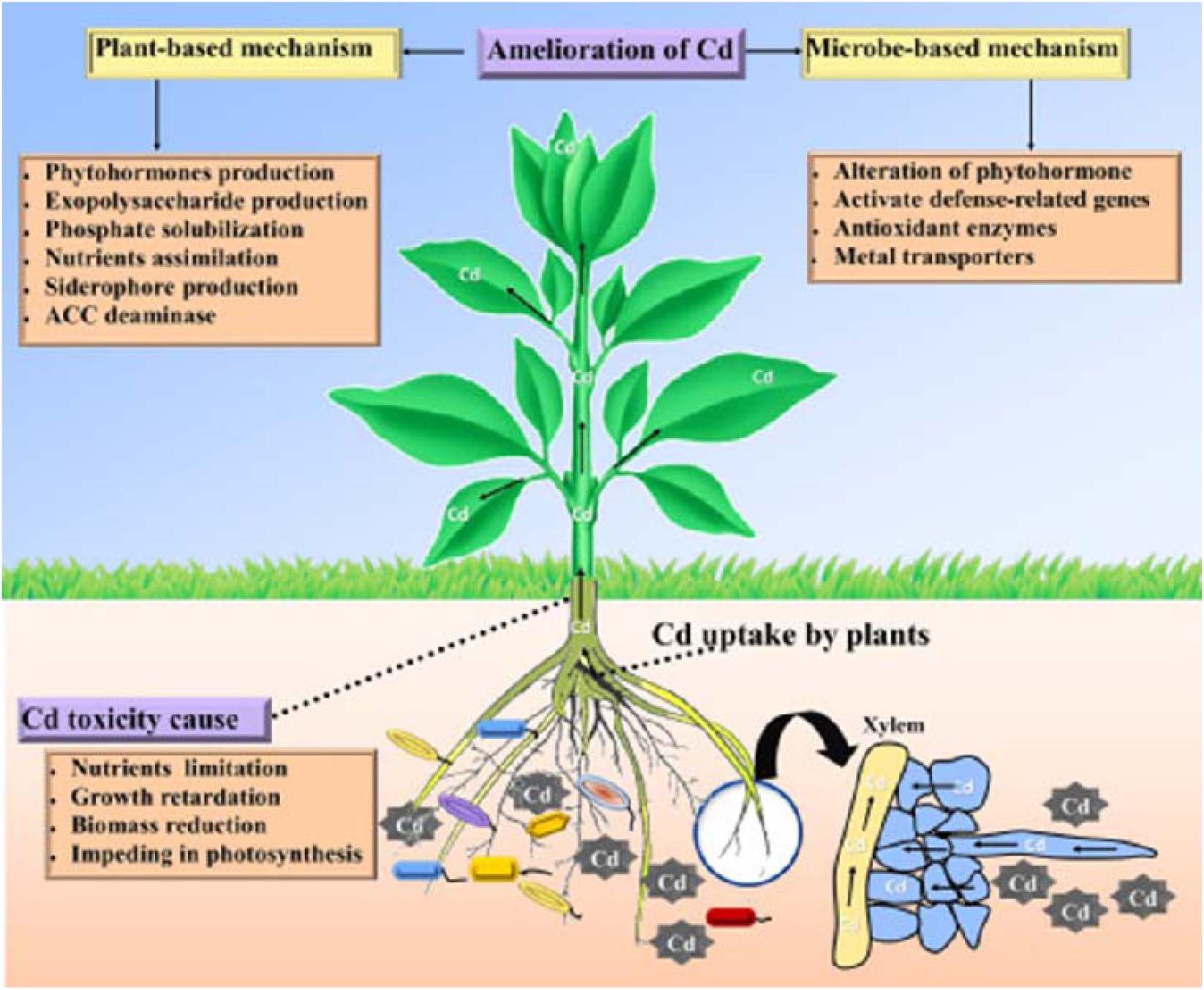
Accumulation and effect of Cd^2+^ ions in plant organisms 30 [Shahid].

Such results prompt the study of the effect of cadmium intoxication on the body of experimental animals.

The studies of calcium content allowed to establish that under conditions of cadmium intoxication in the brain of experimental animals the level of this element significantly decreased on the 14th and 28th day by 5 and 7.8 times, respectively, while in the myocardium a decrease by 3 times on the 14th day and an increase by 2.6 times on the 28th day was observed compared to the control group of animals (Table 1).

**Table 1.** Dynamics of changes in the content of elements in the studied organs and tissues under conditions of cadmium intoxication (M ± m, n=10)

| Groups of animals | Organs and tissues under investigation |  |  |  |
| --- | --- | --- | --- | --- |
|  | Brain | Myocardium | Brain | Myocardium |
|  | Calcium, mg/g ash |  | Magnesium, mg/g ash |  |
| Control | $1,96 \pm 0,31$ | $0,69 \pm 0,12$ | $1,08 \pm 0,15$ | $0,62 \pm 0,08$ |
| 1st day | $2,62 \pm 0,83$ | $0,41 \pm 0,10^*$ | $3,49 \pm 0,94^*$ | $1,26 \pm 0,32^*$ |
| 14th day | $0,37 \pm 0,05^*$ | $0,22 \pm 0,06^*$ | $6,49 \pm 0,74^*$ | $1,88 \pm 0,62^{**}$ |
| 28th day | $0,25 \pm 0,06^*$ | $1,83 \pm 0,14^*$ | $2,51 \pm 0,24$ | $1,67 \pm 0,68^*$ |
|  | Copper, mg/g ash |  | Zinc, mg/g ash |  |
| Control | $0,16 \pm 0,01$ | $0,24 \pm 0,06$ | $0,78 \pm 0,21$ | $0,56 \pm 0,16$ |
| 1st day | $0,18 \pm 0,04$ | $0,32 \pm 0,04$ | $0,36 \pm 0,15^{**}$ | $1,09 \pm 0,29^{**}$ |
| 14th day | $0,17 \pm 0,05$ | $0,22 \pm 0,05$ | $1,73 \pm 0,13^*$ | $0,63 \pm 0,18$ |
| 28th day | $0,12 \pm 0,01^*$ | $0,24 \pm 0,06$ | $0,94 \pm 0,20$ | $1,32 \pm 0,28^*$ |

Note: \* – $p < 0.001$ , \*\* – $p < 0.01$ - reliability of the difference in data compared to the control group of animals.

Calcium metabolism is closely related to magnesium metabolism. Determination of Mg content in the brain and myocardium of experimental animals (Table 1) showed a clear tendency to accumulation of this element: starting from the 1st day, the level of Mg increased in the brain - 3.2 times, in the myocardium - 2 times compared to the control values. The maximum increase of magnesium ions was observed on day 14 in the brain (p<0.001) - 6 times and in the myocardium (p<0.01) - 3 times compared to the control group of animals.

The study of the essential trace element copper allowed an establishment of a decrease of this indicator in the brain (p<0.001) by 20% in the late period of the study compared to the control group. Regarding the level of Cu in the myocardium (Table 1), it is necessary to note an increase in the 1st day of intoxication by 33%, followed by a decrease in its concentration to the level of control animals on the 28th day.

Regarding the concentration of zinc, (p<0.001) a 2-fold increase in the early and late periods in the myocardium compared to the control group of animals has been observed. In the brain on the 1st day (p<0.01) the zinc content decreased by 2 times, on the 14th day (p<0.001) this indicator increased by 2 times, and at the end of the experiment it was within the control group of animals.

When administering cadmium chloride, we found a statistically significant (p<0.001) increase in its concentration in the studied organs and tissues. In particular, starting from the 1st day of observation and until the end of the experiment, the cadmium content increased in the brain (p<0.01) - by 60% on the 14th day, in the myocardium (p<0.001) - by 40-42 times, during the entire study period compared to the control group of animals (Figure 3).

**Figure 3.**
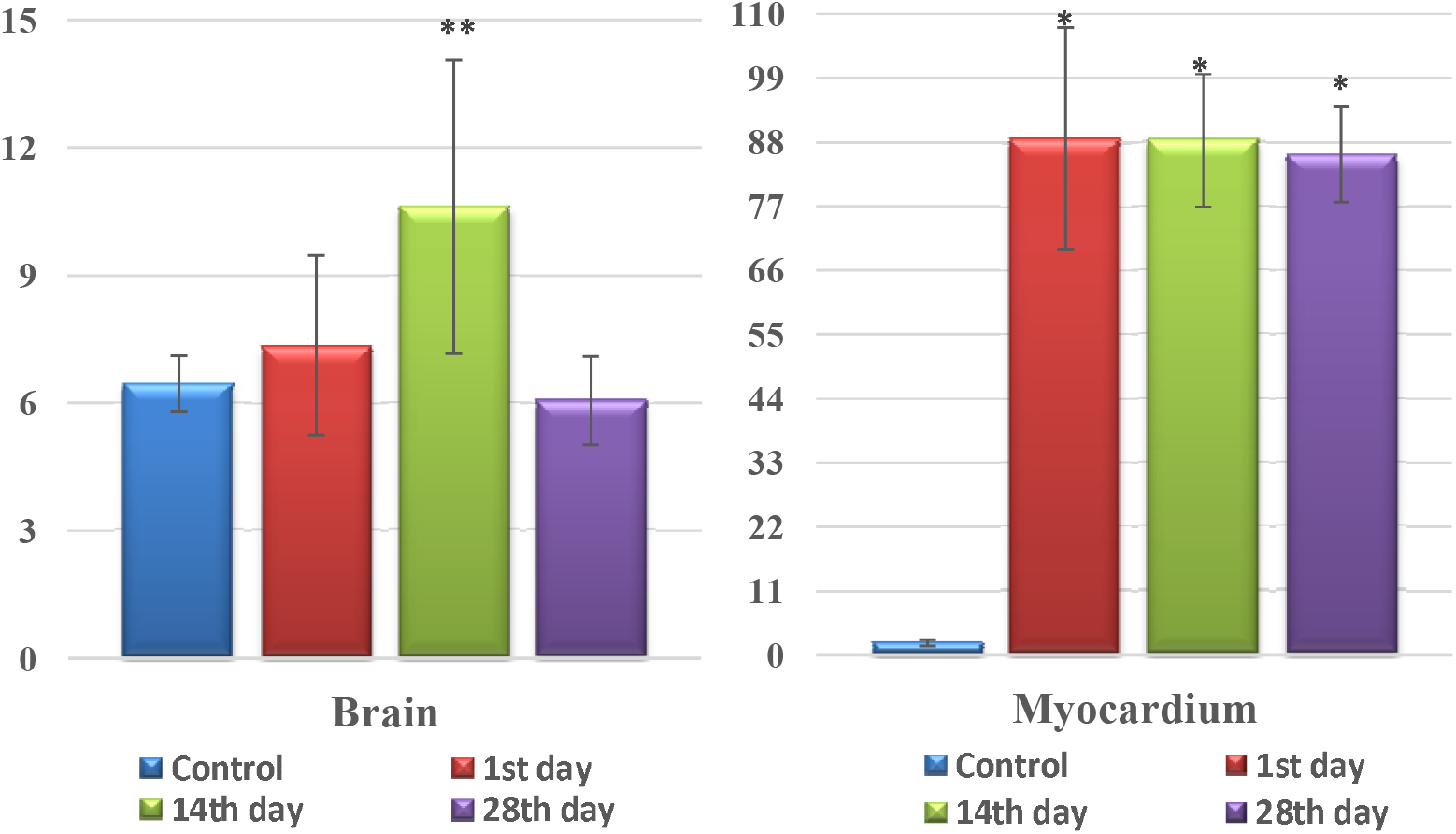
Dynamics of changes in the level of Cadmium in the studied organs and tissues under conditions of cadmium intoxication, μg/g ash

## Discussion

Ensuring environmental stability is possible taking into account all natural and anthropogenic factors that affect the ecological potential of the Prykarpattia region, and subsequently the health of the population. Therefore, it is important to have information on the level of toxic elements, in particular cadmium in soils and water bodies of the region, which allows us to understand their impact on living organisms. In addition, their content in soils and water correlates with their content in biological environments of humans and animals, that is, it may indicate pathological conditions. Soils are the main depositing medium where metals can enter with precipitation and water. Soil conditions should be considered as an integral indicator of the environmental pollution process.

The study of cadmium content in the soils of the region is not accidental, as the authors argue [31,32] that the excessive intake of heavy metals, in particular cadmium, is due to the location of sources of pollution with this metal in the Prykarpattia region, in particular Burshtyn TES; “Oriana” potash plant, Kalush mine; PJSC Ivano-Frankivskcement.

Our soil analysis has shown an increase in cadmium levels in soils of different geographical zones of the region. The increase in cadmium levels in soils can be caused by anthropogenic emissions from enterprises, waste from livestock farms, agricultural activities, the use of pesticides, and vehicle emissions [33, 34].

Along with this, soil pollution is a source of surface and groundwater pollution. Water and salts dissolved in it play an integral role in ensuring the constancy of the internal environment of the body. The content of chemical elements in water depends on many factors, among which the main place belongs to human activity. At the same time, it was found that in the drinking water of the plain and foothill zone the level of cadmium exceeded the maximum permissible level. According to the literature [4, 5, 7], the increase in cadmium levels may be due to the pollution of water bodies by industrial and domestic wastes with organic and mineral fertilizers, especially phosphate fertilizers.

The analysis of drinking water allowed us to determine that a significant number of people living in the plain and foothill zone consume water with a high content of cadmium. It leads to an increase in the effect of this toxicant on living organisms.

Pollution with heavy metal salts reduces plant productivity, disrupts natural phytocoenosis, causes destruction of the assimilative potential of phytomass, leads to disruption of organogenesis processes in the form of specific changes in plants and worsens the quality of agricultural products [35]. It is known from scientific sources [8] that plants growing in soil contaminated with cadmium are the main source of cadmium intake into the human body.

Assessment of cadmium levels in environmental objects and understanding of the mechanisms of its impact on living organisms is crucial for the prevention of a number of pathologies. In particular, the data obtained served as a basis for studying the effect of cadmium salts on the macro- and microelement composition of the body of experimental animals.

It is known from the literature about the antagonistic properties of cadmium ions in relation to such macroelements as calcium and magnesium and essential trace elements zinc and copper. Therefore, it is important to study the distribution of these elements in various organs and tissues in the dynamics of cadmium intoxication.

The obtained results showed that under cadmium intoxication in the body of experimental animals there are changes in the level of macronutrients-calcium and magnesium. In particular, there is a decrease in the level of calcium in the brain on the 14th and 28th day - 5-7.8 times, myocardium on the 1st and 14th day - 3 times, respectively, followed by an increase on the 28th day compared to the control group. At the same time, the level of magnesium in the brain and myocardium significantly increased during the entire observation period.

Analysis of the level of copper showed a decrease in the brain on the 28th day by - 20%; in the myocardium, the level of copper increased on the 1st day by 33%. However, at the end of the experiment it did not exceed the value of control animals.

The study of zinc levels showed that this indicator increased most of all on the 14th day in the brain by 2 times compared to the control values. In the myocardium, the zinc content was higher than in control animals during the entire observation period.

The conducted studies allowed to establish that in animals affected by cadmium chloride there was a significant violation of the level of macroelements such as Ca, Mg, and essential trace elements Zn and Cu.

Such changes in the level of essential macro- and microelements occurred against the background of accumulation of cadmium in these organs. Its level increased in the brain - by 60% on day 14, in the myocardium - by 40-42 times compared to the control group of animals.

The results obtained are consistent with the literature data. In case of violation of physiological levels of metals in tissues and organs, competition for chemical active groups and catalytic centers in macromolecules with changes in their structure and functions is possible [36,37]. On the other hand, changes in the level of macro- and microelements can be considered as an adaptive reaction - the body’s response to the intake of cadmium ions. Cadmium accumulation can be explained by the disruption of metallothionein proteins, which are able to influence the metabolism of this element. Thus, the excessive intake of cadmium salts causes the development of dysmicroelementosis, which is accompanied by a violation of the homeostasis of essential macro- and microelements. At the same time, the disturbance of homeostasis of such elements causes changes in the biochemical functions of many enzymes that contribute to the initiation of cascade reactions, which leads to the development of oxidative stress, impaired energy supply, signal transduction, etc.

## Conclusions

1. Significant differences in the content of cadmium in soils and water of the plain, foothill and mountain zones of the Prykarpattia region were established. In particular, an increase of cadmium in soils of plain and mountainous zones of the region has been noted.
2. The increase of cadmium concentration in water exceeding the maximum permitted levels is observed in plain and foothill areas with intensive development of agriculture and industry, and the minimum level is in mountainous areas.
3. The literature data on the main ways of supply and accumulation of heavy metal ions, in particular cadmium, have been analyzed. It has been established that cadmium accumulates mainly in the root system of plants and partially in the aerial part. Their negative impact on the plant organism has been described.
4. Significant disorders in the body of experimental animals under conditions of excessive intake of cadmium ions have been revealed. It was accompanied by the accumulation of cadmium in the myocardium and brain, against the background of redistribution of vital macronutrients calcium and magnesium and micro-elements copper and zinc. Thus, excessive intake of cadmium salts causes the development of dysmicroelementosis, which is accompanied by a violation of the homeostasis of a living organism.
5. Continuous monitoring of toxicants in the ecosystem is recommended as an important component of environmental monitoring.

This issue requires further research to determine the level of toxicants in the environment. Furthermore, it is also a prerequisite for further experimental studies on the impact of cadmium compounds on the micro-element status of the body and ways to protect against such disorders.

## Notes

### Competing Interest Statement

The authors have declared no competing interest.

